# Modeling the assessment of the upper limb motor function impairment in children with cerebral palsy using sEMG and IMU sensors

**DOI:** 10.1101/748202

**Authors:** S. Raouafi, M. Raison, A. Sofiane

**Author notes:** Corresponding author: CRME – Research Center, Office GR-123, 5200, East Bélanger Street, H1T 1C9, Montréal, QC, Canada, # 8189.

## Abstract

Several rehabilitation approaches have shown that robot-assisted therapy (robot-AT) can improve the quality of upper limb movements in children with cerebral palsy (CP). However, there is still no method for assessing upper limb motor function impairment using a combination of surface electromyography (sEMG) and inertial measurement unit (IMU) sensors. The aim of this study was to develop a functional ability model to assess the effectiveness of robot-AT on improving upper limb function in children with CP. Fifteen healthy children and fifteen children with CP were included in this study. Children with CP performed eighteen robot-AT sessions and were evaluated twice, using EMG and three-axis IMU readings from accelerometer (IMU-ACC). Principal component analysis and the RELIEFF algorithm were used for dimensionality reduction of the feature space. The classification was performed by using support vector machines, linear discriminant analysis, and random forest. The proposed assessment method was evaluated by using leave-one-out cross validation. With this approach, it was possible to differentiate between healthy children and children with CP pre-robot-AT and post-robot-AT with an overall accuracy of 97.56%. This study suggests that there is potential for modeling the assessment of the upper limb motor function impairment in children with CP using sEMG and IMU-ACC sensors.

## 1. Introduction

Cerebral palsy (CP) is the most common motor disability in childhood, affecting approximately 2 to 2.5 per 1000 children born (Bax et al., 2005). It is characterized by low muscle tone and poor movement coordination. This disability has a major negative and devastating effect on children’s development and on their quality of life. About 50% of children with CP present loss of functions of their upper extremities. Consequently, in the past decades, the focus of research in this field has been on the assessment of upper limb motor function, which is crucial for clinicians to use during CP therapy to evaluate the patient rehabilitation needs and increase the chances of a prognosis of motor function improvement. Consequently, this would increase the autonomy of CP children. In clinical practice, different tools were developed to evaluate UL function for individuals with CP (Eliasson et al., 2006; Fedrizzi, Pagliano, Andreucci, Oleari, & neurology, 2003; House, Gwathmey, Fidler, & volume, 1981). Among all these methods, the MACS is considered the most reliable and valid tool to evaluate performance of upper limb tasks in daily living (Plasschaert, Ketelaar, Nijnuis, Enkelaar, & Gorter, 2009).

While some of these tools have been validated, the current outcome measures are mainly qualitative and therefore subjected to the perspective of the therapist evaluating the child. Moreover, these tools do not consider pathological mechanisms, which make it more difficult for therapists aiming to assess the effectiveness of a specific intervention (Gajewska, Sobieska, & Samborski, 2006).

Recent studies have shown the potential of quantitative electromyography (EMG) analysis in the identification of gait patterns in patients with cerebral palsy. For example (Sangeux, Rodda, & Graham, 2015) used a dataset of 776 CP patients, i.e. 1552 limbs, to compare their sagittal gait patterns using K-means clustering. EMG parameters have also been found to help distinguish between healthy subjects and patients with muscle disorders like Duchenne muscular (Hogrel, 2005). (Abel, Zacharia, Forster, Farrow, & physics, 1996)developed different learning algorithms for EMG-based diagnosis of neuromuscular disorders. In this study, results showed that EMG segment analysis achieved 60–80% accuracy in classifying healthy, myopathic and neuropathic subjects. In a recent study, EMG signals were used to generate a visual biofeedback signal for wrist movement rehabilitation. It was concluded that Shannon entropy could provide an accurate visual biofeedback for reduction of spasticity in patients with a stroke (Zadnia, Kobravi, Sheikh, Hosseini, & Neuroscience, 2018). Another research study found that implementing an EMG-based model of muscle health in a rehabilitative elbow brace has the potential of assessing patients recovering from Musculoskeletal (MSK) elbow trauma (Farago, 2018). Existing methodologies in pattern recognition suggest that EMG signals could be analyzed to detect muscle strength and motor unit (MU) recruitment as well as measure the effectiveness of rehabilitation therapies. Specifically, for children with CP, EMG analysis has been recognized as a robust tool for identifying gait patterns(Gopura, Bandara, Gunasekara, & Jayawardane, 2013).

To the best of our knowledge, only a limited number of studies have attempted to develop a statistical classification method to assess and characterize disability levels(Li et al., 2017). Similarly, only a few clinical studies have attempted to study the effectiveness of robot-assisted therapy (Robot-AT) for children with CP(Chen & Howard, 2016). In these research studies improvements in upper limb function were often assessed based on the opinion of the evaluating therapist. In this context, an accurate EMG-based function ability model could be a helpful tool to better understand movement limitations in terms of muscular behavior and assess the effectiveness of assistive technology interventions for CP children.

The primary objective of the current study is to develop an EMG-based functional ability model to differentiate between healthy and CP children with an appropriate number of EMG features. A secondary objective is to evaluate the effectiveness of robot-AT for children with CP and to examine the added value of an Inertial Measurement Unit-Accelerometer (IMU-ACC) for identifying disability levels.

## 2. Materials and Methods

### 2.1. Participants

Fifteen patients (mean age: 9 years 5 months, SD: 3 years 1 month) were recruited from a school for children with physical disabilities (École Victor-Doré, Montréal, Qc, Canada). Furthermore, fifteen typically developing (TD) children were included as a reference group (mean age: 8 years 2 months, SD: 2 years 7 months). This sample size was dependent on the recruitment capacity of the school. The patients’ characteristics are described in Table 1. The inclusion criteria included a history of CP, a maximum age of 14 years and the ability to understand or perform the tasks. Among the exclusion criteria were botulinum toxin injections within six months before measurements or previous orthopedic surgery in the upper limbs. The study was approved by the Research Ethics Boards of Ste. Justine Hospital. The study entailed the free participation and authorization form was approved by the parents or guardians of the children involved.

**Table 1.**
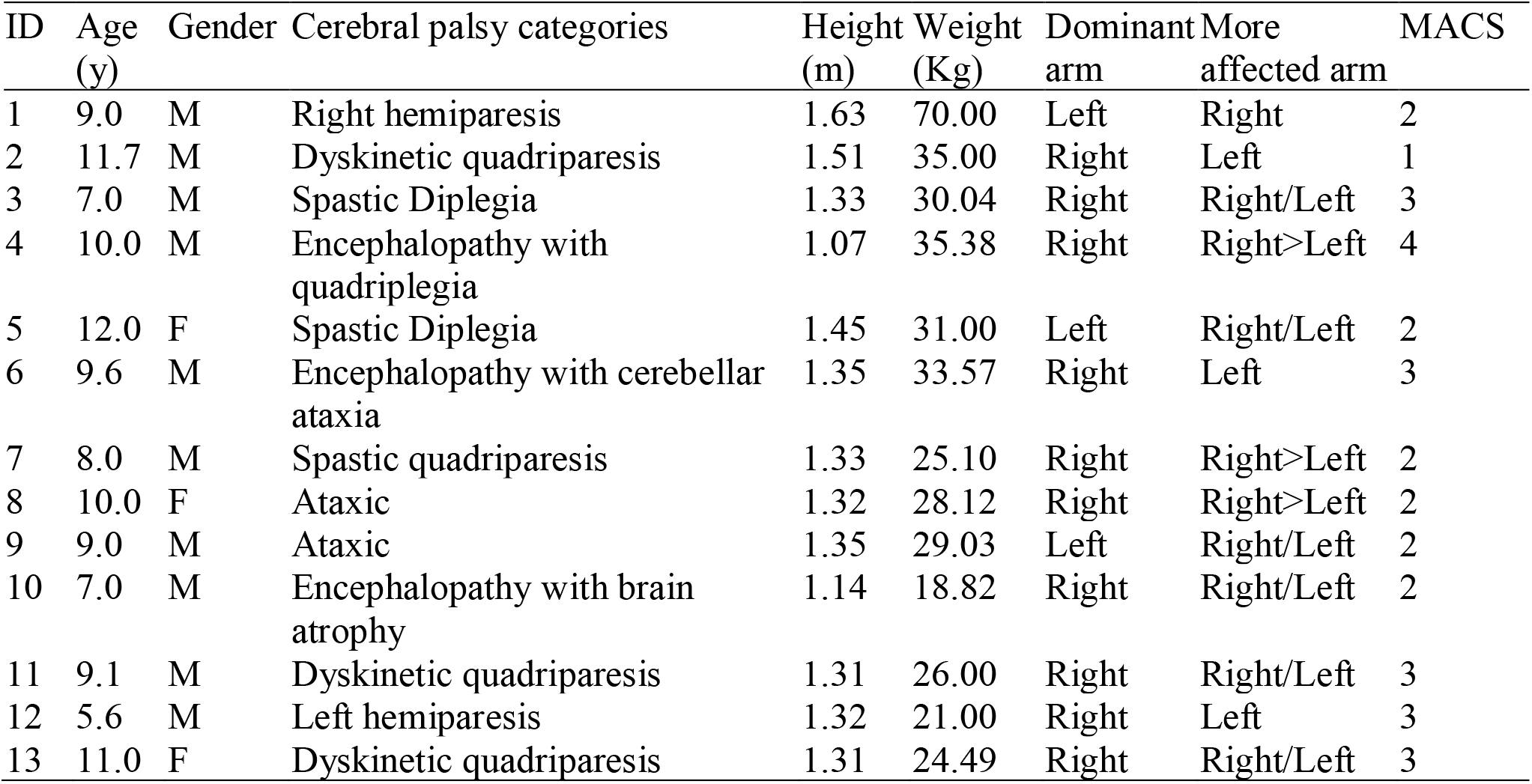
Patient’ characteristics

| ID | Age<br>(y) | Gender | Cerebral palsy categories | Height<br>(m) | Weight<br>(Kg) | Dominant<br>arm | More<br>affected arm | MACS |
| --- | --- | --- | --- | --- | --- | --- | --- | --- |
| 1 | 9.0 | M | Right hemiparesis | 1.63 | 70.00 | Left | Right | 2 |
| 2 | 11.7 | M | Dyskinetic quadriparesis | 1.51 | 35.00 | Right | Left | 1 |
| 3 | 7.0 | M | Spastic Diplegia | 1.33 | 30.04 | Right | Right/Left | 3 |
| 4 | 10.0 | M | Encephalopathy with<br>quadriplegia | 1.07 | 35.38 | Right | Right>Left | 4 |
| 5 | 12.0 | F | Spastic Diplegia | 1.45 | 31.00 | Left | Right/Left | 2 |
| 6 | 9.6 | M | Encephalopathy with cerebellar<br>ataxia | 1.35 | 33.57 | Right | Left | 3 |
| 7 | 8.0 | M | Spastic quadriparesis | 1.33 | 25.10 | Right | Right>Left | 2 |
| 8 | 10.0 | F | Ataxic | 1.32 | 28.12 | Right | Right>Left | 2 |
| 9 | 9.0 | M | Ataxic | 1.35 | 29.03 | Left | Right/Left | 2 |
| 10 | 7.0 | M | Encephalopathy with brain<br>atrophy | 1.14 | 18.82 | Right | Right/Left | 2 |
| 11 | 9.1 | M | Dyskinetic quadriparesis | 1.31 | 26.00 | Right | Right/Left | 3 |
| 12 | 5.6 | M | Left hemiparesis | 1.32 | 21.00 | Right | Left | 3 |
| 13 | 11.0 | F | Dyskinetic quadriparesis | 1.31 | 24.49 | Right | Right/Left | 3 |

### 2.2. Data acquisition

Children with CP performed two robot-AT sessions with the REAplan end-effector rehabilitation robot (Gilliaux et al., 2015) and two occupational therapy sessions per week over a period of ten weeks. Each session lasted approximately 45 minutes. The children were evaluated twice, before and after each intervention (i.e. on week 1 and week 10). Each child was assessed using the REAplan by moving the upper limb in a horizontal plane (Sapin, Dehez, Vanderwegen, Gilliaux, & Lejeune, 2013).

MACS scores were used to evaluate the children upper limb manual capacity(Eliasson et al., 2006). Healthy children were evaluated once with the REAplan. During each evaluation session, each child was comfortably seated, and the upper limb fixed on the end effector of the robotic device. Movement performance was computed from two unidirectional tasks (a target task and a free Amplitude task) and two geometrical tasks (circle task and a square task). Each child was instructed to perform the four tasks ten times. sEMG and IMU-ACC data were collected from wearable sensors (Trigno, Delsys Inc., Natick, MA, USA). Each sensor included information from both muscle activation, via surface EMG sensors, and position by triaxial ACC. Eight sensors were attached to the skin surface of the anterior deltoid, lateral deltoid, posterior deltoid, biceps brachii, triceps Brachii, infraspinatus, brachialis and brachioradialis (Fig. 1), according to the guidelines of the SENIAM project (Hermens, Freriks, Disselhorst-Klug, & Rau, 2002). EMG signals were pre-amplified and band pass filtered (from 20 to 450Hz) with a sampling frequency of 1111.11 Hz. The accelerometer sampling frequency was 148.1 Hz.

**Figure 1.**
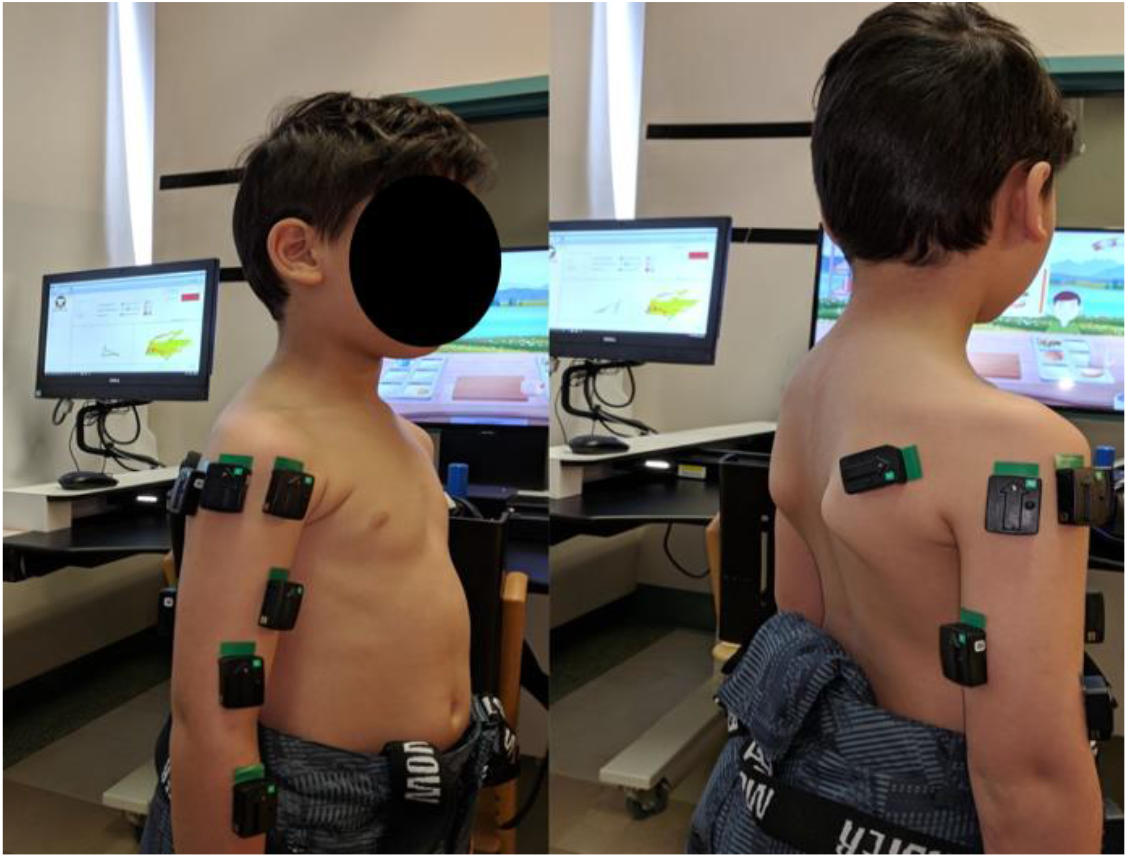
Placement of the eight sensors

### 2.3. Data processing

The sEMG and IMU-ACC data were processed using custom-written MATLAB® (MathWorks®, Natick, USA) scripts.

#### Post-trial selection criteria

Datasets from two subjects were excluded from further analysis due to: 1. Data corrupted after visual inspection; 2. Hardware failure during the data acquisition.

#### Feature extraction

The most widely used features from time and frequency domains, as described in the literature, were extracted from each EMG segment (Appendix 1). These chosen features are commonly used for pattern recognition and real-time applications (Phinyomark, Phukpattaranont, & Limsakul, 2012; Scheme, Englehart, & Development, 2011).

In addition, six features were extracted from the magnitude of the acceleration including the mean, the standard deviation, the maximum, the minimum, the local minimum and maximum peaks.

#### Feature reduction

First, the best individual features were selected by comparing their individual performances using majority vote models on the four tasks (James, 1998). Feature sets were then developed. In order to further optimize feature sets, dimensionality reduction was performed. Two dimensionality-reduction techniques were used, principal component analysis (PCA) (Dallas, 2014) and the RELIEFF algorithm.

### 2.4. Classification

Each model was evaluated for each of the four tasks separately. Three classification models were investigated: The Linear discriminant Analysis(James, Witten, Hastie, & Tibshirani, 2013), the Support Vector Machine (SVM) with polynomial kernel function and one-vs-one scheme (James et al., 2013), and the Random Forest (Patel et al., 2010). Classification models were developed to differentiate between two and three classes. First, we created LDA, SVM, and RF classifiers to distinguish between healthy children and children with CP. Secondly, the same classifiers were used to distinguish between healthy children and children with CP pre_robot-AT and post_robot-AT, respectively. Since we used different types of features, a classifier-level fusion was performed to combine results from classifiers trained separately by sEMG and IMU-ACC feature subsets. Vectors of posterior probabilities of the two first-level classifiers were concatenated to train the final model.

In the two-class models, the classification accuracy was measured using a leave-one-out patient cross-validation (LOOCV) (Kazerouni, Achiche, Hisarciklilar, & Thomson, 2011; Wen, Raison, & Achiche, 2018). In the three-class models, datasets from healthy children and children with CP pre_robot-AT and post_robot-AT were randomly split into approximately 70% for training and 30% for testing each. This was done in order to handle dataset imbalances. The process was then repeated 10 times. The 10 values of accuracy were averaged to produce the final accuracy.

## 3. Results

The individual performances for each feature after applying a majority vote among the four tasks have been included as supplementary material (Appendix 2). Models were evaluated using a LOOCV. The RF classifier provided the highest accuracy for most sEMG features. New feature subsets were developed. Feature subset 1 (FS1) included only Mean Spike Slope (MSS) and Multiple Window (MHW) features (with 85.71 % and 82.14% accuracy respectively), because adding more features caused the degradation of the classification performance. Feature subset 2 (FS2) included the top ranked features among each category of sEMG features, these are: Spectral Moment 2 (SM2), Integrated EMG (IEMG), Average Amplitude Change (AAC), Zero Crossing (ZC), MHW, Coefficients of Cepstral Analysis (CC4), Maximum Fractal Length (MFL) and MSS. Skewness (SKEW), and Shannon entropy (SHANNON) were excluded because of the low classification accuracy. The IEMG, MHW, and CC4 features were selected as the best features for all tasks with RELIEFF algorithm and were considered as Feature subset 3 (FS3). For accelerometric signals, LDA provided the highest accuracy among all tasks and only the mean feature (89.26% accuracy) was considered for further analysis. First, models were developed to classify between the healthy and CP children pre-Robot-AT. FS2 provided better classification accuracy when used with the RF classifier (67.86–85.71%) than those achieved with the SVM classifier (46.42-53.57%). Classification accuracies did not improve after dimensionality reduction. Performances were ranged between 39.29 and 75% with RELIEFF algorithm and 46.43-71.43% with PCA. For IMU-ACC data, mean feature tended to work better with the LDA classifier and provided the best accuracies among all tasks (82.14-92.86%). PCA did not improve classifier performances and reached a maximum accuracy of 71.43%. All classification accuracies for sEMG and IMU-ACC feature subsets are presented in Table 2.

**Table 2.** Classification accuracies for all feature subsets (two-class models)

| Classification accuracy (%) |  |  |  |  |
| --- | --- | --- | --- | --- |
| Feature subset | Task | LDA | SVM | RF |
| FS1 | Square | 60.71 | 53.57 | **82.14** |
|  | Free amplitude | 64.29 | 57.14 | 75.00 |
|  | Circle | 53.57 | 53.57 | 75.00 |
|  | Target | 60.71 | 60.71 | 75.00 |
| FS2 | Square | 64.29 | 46.42 | 75.00 |
|  | Free amplitude | 46.43 | 46.42 | 67.86 |
|  | Circle | 53.57 | 46.42 | 82.14 |
|  | Target | 32.14 | 46.42 | **85.71** |
| FS3 (RELIEFF) | Square | 53.57 | 50.00 | **75.00** |
|  | Free amplitude | 46.43 | 57.14 | 71.43 |
|  | Circle | 60.71 | 39.29 | **75.00** |
|  | Target | 42.86 | 67.86 | **75.00** |
| F4 (PCA) | Square | 60.71 | 46.43 | 50.00 |
|  | Free amplitude | 57.14 | **71.43** | 53.57 |
|  | Circle | 46.43 | 46.43 | 64.29 |
|  | Target | 67.86 | 46.43 | **71.43** |
| FS1-IMU-ACC | Square | **92.86** | **92.86** | 89.29 |
|  | Free amplitude | 82.14 | 78.57 | 78.57 |
|  | Circle | **92.86** | 89.29 | 89.29 |
|  | Target | 85.71 | 85.71 | 82.14 |
| FS2-IMU-ACC<br>(PCA) | Square | 50.00 | 46.43 | 53.57 |
|  | Free amplitude | 57.14 | 53.57 | **71.43** |
|  | Circle | 53.57 | 46.43 | 60.71 |
|  | Target | 53.57 | 46.43 | **71.43** |

Classification accuracies for fused data from sEMG and IMU-ACC were the same for all tasks. Final classifier inputs were obtained from trained RF classifier for sEMG features and LDA classifier for IMU-ACC features. SVM classifier provides the best accuracy for all tasks with fused data as inputs (96.43%).

Following the development of two-class models (Healthy as class1 and CP children pre-robot-AT as class2), classifiers were developed to distinguish between three categories: healthy, CP children pre-robot-AT and CP children post. Results are shown in Table 3.

**Table 3.** Classification accuracies for all feature subsets (Three-class models)

| **Classification accuracy** |  |  |  |  |
| --- | --- | --- | --- | --- |
| **Feature subset** | **Motion** | **LDA** | **SVM** | **RF** |
| FS1 | Square | 35.83 | 33.33 | 48.33 |
|  | Free amplitude | 41.67 | 33.33 | 51.67 |
|  | Circle | 38.33 | 33.33 | 47.50 |
|  | Target | 39.17 | 33.33 | 50.83 |
| FS2 | Square | 32.50 | 38.33 | 49.17 |
|  | Free amplitude | 43.33 | 59.17 | 50.83 |
|  | Circle | 43.33 | 55.00 | 55.83 |
|  | Target | 50.83 | 44.17 | 54.17 |
| FS3 (RELIEFF) | Square | 33.33 | 33.33 | 51.67 |
|  | Free amplitude | 31.67 | 33.33 | 43.33 |
|  | Circle | 48.33 | 33.33 | **61.67** |
|  | Target | 51.67 | 33.33 | 55.83 |
| F4 (PCA) | Square | 35.00 | 33.33 | 58.33 |
|  | Free amplitude | 40.00 | 35.83 | 44.17 |
|  | Circle | 35.83 | 41.67 | 51.67 |
|  | Target | 46.67 | 33.33 | 42.50 |
| FS1-IMU-ACC | Square | 63.33 | 74.17 | 73.33 |
|  | Free amplitude | 57.50 | 57.50 | 72.50 |
|  | Circle | 62.50 | **78.33** | 73.33 |
|  | Target | 65.83 | 70.83 | 69.17 |
| FS2-IMU-ACC (PCA) | Square | 27.50 | 33.33 | 42.50 |
|  | Free amplitude | 38.33 | 30.00 | 51.67 |
|  | Circle | 35.83 | 33.33 | 37.50 |
|  | Target | 40.83 | 33.33 | 37.50 |

FS3 provided better classification accuracy when used with the RF classifier (43.33–61.67%) than those achieved with the SVM and LDA classifiers (33.33% for SVM and 33.33-51.67% for LDA). Classification accuracies remain low after dimensionality reduction with PCA (33.33-58.33%). For IMU-ACC feature, SVM provided the best accuracy among all tasks (57.50-78.33%). Regarding the final classification, fused data improved classification performances to achieve 97.56% for circle task using RF model (Fig .2).

**Figure 2.**
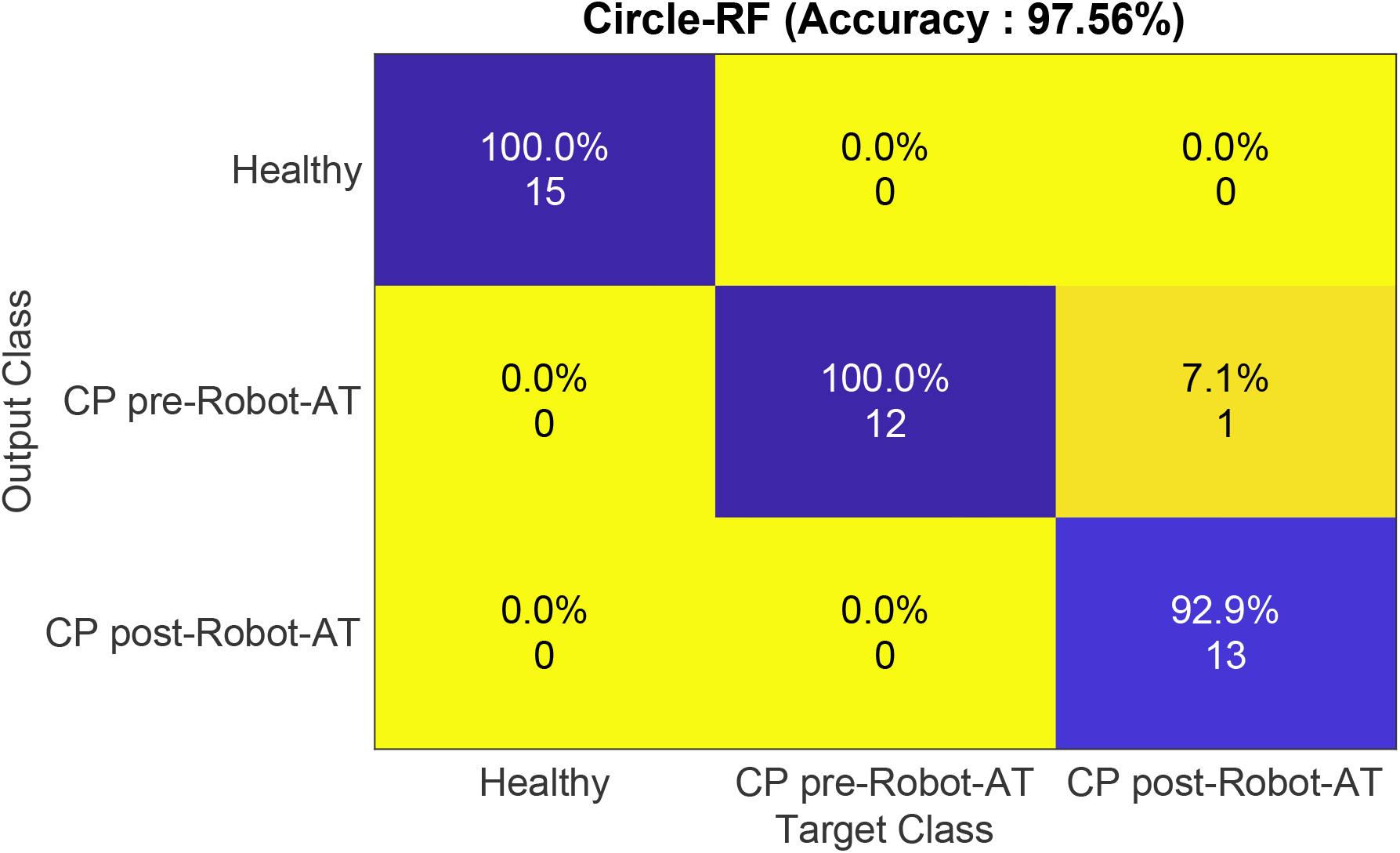
Interposition classification accuracies, averaged across all subjects

## 4. Discussion

Classification techniques were used in this study to identify disability levels of children with CP after Robot-AT. The purpose of this classification was to assess the effectiveness of Robot-AT and to provide support for clinical evaluation in classifying children based on features extracted from EMG and IMU-ACC signals. The main result of this study was that SVM classifier performed well in distinguishing between healthy individuals and those with cerebral palsy based on fused data from sEMG and IMU-ACC signals (97.56% classification rate was obtained).

For two-class models, confusion matrix showed that one patient disabled at level I, according to MACS, was misclassified. Further, the confusion matrix of the three-class model (Fig.2,) showed that one patient in the pre-Robot-AT evaluation was misclassified. The diagnosis of this latter patient was spastic quadriparesis and, on the basis of magnetic resonance imaging (MRI), a diagnosis of intramedullary tumor. According to MACS, this patient is considered to have a disability level 2 pre-Robot-AT and a minor functional limitation. This suggests that classification accuracy might be limited for patients with mild to moderate motor dysfunction. It should be reminded that all tasks in the evaluation process are made in a small workspace with a standard and circular manner without regard to patient’s specific characteristics and abilities. In this context, results may be misleading, and patient misclassified as healthy or with a minor functional limitation. To overcome this, a patient-specific evaluation tasks and an individualized trajectory should be designed, depending upon each patient’s type of injury, disability level, range of motion and performance.

Our results showed that adding IMU-ACC features to the optimal feature subset (IEMG, MHW, and coefficients of spectral analysis) produced an average increase of 35.89% in the classification accuracy. The results suggest that these features are more efficient to identify muscle health than the more commonly used sEMG features. This is in accordance with (Abel et al., 1996; Farago, 2018; Haddara, 2016; Hogrel, 2005), who reported that features extracted from sEMG can distinguish between normal and abnormal muscle patterns.

The proposed assessment method can serve as a potential approach to assess the effectiveness of Robot-AT and of other upper limb conventional therapies. Moreover, this method can also be used as part of an adaptive algorithm for the REAplan.

One limitation of this approach, however, is the small sample size and thus larger study cohorts are required to validate the proposed EMG-IMU-ACC based functional ability model to assess the effectiveness of Robot-AT for children with CP.

## Acknowledgments

We gratefully acknowledge the contribution of the Canada Foundation for Innovation (CFI). The authors would like to thank the patients who participated in this study as well as the occupational therapist’s team at “École Victor-Doré” for their collaboration.

Supporting Information (Appendix 1 and Appendix 2) are available online.

## Abbreviations

ACC: Accelerometer
CP: Cerebral Palsy
EMG: Electromyography
FS: Feature subset
IMU: Inertial Measurement Unit
LDA: Linear Discriminant Analysis
MSK: Musculoskeletal
PCA: Principal Component Analysis
Robot-AT: Robot-Assisted Therapy
RF: Random Forest
SVM: Support Vector Machine

## References

Abel, E., Zacharia, P., Forster, A., Farrow, T. J. M. e., & physics. (1996). Neural network analysis of the EMG interference pattern. 18(1), 12–17.

Bax, M., Goldstein, M., Rosenbaum, P., Leviton, A., Paneth, N., Dan, B., … neurology, c. (2005). Proposed definition and classification of cerebral palsy, April 2005. 47(8), 571–576.

Chen, Y.-P., & Howard, A. M. J. D. n. (2016). Effects of robotic therapy on upper-extremity function in children with cerebral palsy: a systematic review. 19(1), 64–71.

Dallas, G. J. L. a. (2014). Principal Component Analysis 4 Dummies: Eigenvectors, Eigenvalues and Dimension Reduction. 10.

Eliasson, A. C., Krumlinde-Sundholm, L., Rosblad, B., Beckung, E., Arner, M., Ohrvall, A.M., & Rosenbaum, P. (2006). The Manual Ability Classification System (MACS) for children with cerebral palsy: scale development and evidence of validity and reliability. Dev Med Child Neurol, 48(7), 549–554. doi:10.1017/S0012162206001162

Farago, E. (2018). Development of an EMG-based Muscle Health Model for Elbow Trauma Patients.

Fedrizzi, E., Pagliano, E., Andreucci, E., Oleari, G. J. D. m., & neurology, c. (2003). Hand function in children with hemiplegic cerebral palsy: prospective follow-up and functional outcome in adolescence. 45(2), 85–91.

Gajewska, E., Sobieska, M., & Samborski, W. J. C. n. r. i. o. p. (2006). Manual ability classification system for children with cerebral palsy. 71(4), 317–319.

Gilliaux, M., Renders, A., Dispa, D., Holvoet, D., Sapin, J., Dehez, B., … repair, n. (2015). Upper limb robot-assisted therapy in cerebral palsy: a single-blind randomized controlled trial. 29(2), 183–192.

Gopura, R., Bandara, D., Gunasekara, J., & Jayawardane, T. (2013). Recent trends in EMG-Based control methods for assistive robots. In Electrodiagnosis in new frontiers of clinical research: IntechOpen.

Hermens, H., Freriks, B., Disselhorst-Klug, C., & Rau, G. (2002). The SENIAM project: Surface electromyography for non-invasive assessment of muscle. Paper presented at the ISEK Congress.

Hogrel, J.-Y. J. N. C. C. N. (2005). Clinical applications of surface electromyography in neuromuscular disorders. 35(2-3), 59–71.

House, J. H., Gwathmey, F., Fidler, M. J. T. J. o. b., & volume, j. s. A. (1981). A dynamic approach to the thumb-in palm deformity in cerebral palsy. 63(2), 216–225.

James, G. (1998). Majority vote classifiers: theory and applications. Stanford University,

James, G., Witten, D., Hastie, T., & Tibshirani, R. (2013). An introduction to statistical learning (Vol. 112): Springer.

Kazerouni, A. M., Achiche, S., Hisarciklilar, O., & Thomson, V. J. J. o. M. D. (2011). Appraisal of new product development success indicators in the aerospace industry. 133(10), 101013.

Li, Y., Zhang, X., Gong, Y., Cheng, Y., Gao, X., & Chen, X. J. S. (2017). Motor function evaluation of hemiplegic upper-extremities using data fusion from wearable inertial and surface EMG sensors. 17(3), 582.

Patel, S., Hughes, R., Hester, T., Stein, J., Akay, M., Dy, J., & Bonato, P. (2010). Tracking motor recovery in stroke survivors undergoing rehabilitation using wearable technology. Paper presented at the Engineering in Medicine and Biology Society (EMBC), 2010 Annual International Conference of the IEEE.

Phinyomark, A., Phukpattaranont, P., & Limsakul, C. J. E. S. w. A. (2012). Feature reduction and selection for EMG signal classification. 39(8), 7420–7431.

Plasschaert, V., Ketelaar, M., Nijnuis, M., Enkelaar, L., & Gorter, J. J. C. r. (2009). Classification of manual abilities in children with cerebral palsy under 5 years of age: how reliable is the Manual Ability Classification System?, 23(2), 164–170.

Sangeux, M., Rodda, J., & Graham, H. K. (2015). Sagittal gait patterns in cerebral palsy: the plantarflexor-knee extension couple index. Gait Posture, 41(2), 586–591. doi:10.1016/j.gaitpost.2014.12.019

Sapin, J., Dehez, B., Vanderwegen, M., Gilliaux, M., & Lejeune, T. (2013). Développement du robot de rééducation du membre supérieur REAplan. Paper presented at the XLIèmes Journées d’étude de la SORNEST. Technologies innovantes en Médecine Physique et Réadaptation.

Scheme, E., Englehart, K. J. J. o. R. R., & Development. (2011). Electromyogram pattern recognition for control of powered upper-limb prostheses: state of the art and challenges for clinical use. 48(6).

Wen, J., Raison, M., & Achiche, S. J. J. o. B. (2018). Using a cost function based on kinematics and electromyographic data to quantify muscle forces. 80, 151–158.

Zadnia, A., Kobravi, H. R., Sheikh, M., Hosseini, H. A. J. B., & Neuroscience, C. (2018). Generating the Visual Biofeedback Signals Applicable to Reduction of Wrist Spasticity: A Pilot Study on Stroke Patients. 9(1), 15–26.

