## Appendix 1 for "Modeling the assessment of the upper limb motor function impairment in children with cerebral palsy using sEMG and IMU sensors"

Extracted sEMG features from all domains

| **Features** | **Mathematical equation** |
| --- | --- |
| Mean Frequency (*MNF*) [Hz] | $MNF=$ $\frac{\sum_{k=1}^{N} k.S_{x}\left( k \right)}{\sum_{K=1}^{N} S_{x}\left( k \right)}$ |
| Median Frequency (*MDF*) [Hz] | *MDF*= $\frac{1}{2} \sum_{k=1}^{N} S_{x}\left( k \right)$ |
| Peak frequency (*PKF*) [Hz] | $PKF$= (arg (max ($S_{x}\left( k \right)$))  k = 1,…,N |
| Mean Power (*MNP*) [W] | *MNP* _=_ $\frac{\sum_{k=1}^{N} S_{x}\left( k \right)}{M}$ |
| 1^st^, 2^nd^ and 3^rd^ spectral moment [$\mathrm{mV}^{i}$] | ${SM}_{i}$= $\sum_{k=1}^{N} P_{k}f_{k}^{i}$  where $P_{k}$ is the EMG power spectrum at bin k,  $f_{k}$ is the frequency of the spectrum at bin k and  i is the order of the spectral moment. |
| Variance of Central Frequency (*VCF*) [Hz] | *VCF*=$\frac{SM2}{SM0}-\left( \frac{SM1}{SM0} \right)^{2}$ |
| Frequency Ratio (*FR*) | *FR* = $\frac{\sum_{k=f_{1}}^{f_{2}} S_{x}\left( k \right)}{\sum_{f_{1}}^{f_{2}} S_{x} \left( k \right)}$ |
| Integrated EMG (*IEMG*) [mV.s] | $IEMG=\sum_{n=1}^{N} \left\vert x(n) \right\vert$ |
| Root Mean Square (*RMS*) [mV] | $RMS=\sqrt{\frac{1}{N}}\sum_{n=1}^{N} (x(n)^{2})$ |
| Mean Absolute Value (*MAV*) [mV] | $MAV=\frac{1}{N}\sum_{n=1}^{N} \left\vert x(n) \right\vert$ |
| Zero Crossing (*ZC*) | $\left\{ x\left( i \right)>0 and x\left( i+1 \right)<0 \right\}$  Or  $\left\{ x\left( i \right)<0 and x\left( i+1 \right)>0 \right\}$ |
| Waveform Length (*WL*) [mV] | $WL=\sum_{i=1}^{N} \left\vert\Delta x(i) \right\vert$  Δx(i) = x(i) – x(i-1) |
| Willison Amplitude (*WAMP*) | $WAMP=\sum_{i=1}^{N-1} \left[ f(\left\vert x_{i}-x_{i+1} \right\vert) \right]$  Where$\left\{ f\left( x \right)=\left\{ \begin{aligned} 1, &x\geq\varepsilon\\ 0, &\mathrm{otherwise} \end{aligned} \right. \right\}$ |
| Variance of EMG (*VAR*) [mV] | $VAR=\frac{1}{N}\sum_{i=1}^{N} \left( x\left( i \right)- \mu\right)^{2}$  µ=mean value |
| Log-detector (*LOG*) [mV] | *LOG*=$e^{\frac{\sum_{i=1}^{N} \log_{10} \left( \left\vert\left( x_{i} \right) \right\vert\right)}{N}}$ |
| Slope Sign Changes (*SSC*) | $SSC=\sum_{i=1}^{N} f\left[ \begin{aligned} \left( x\left( i \right)-x\left( i-1 \right) \right)\times\\ (x\left( i \right)-x\left( i+1 \right)) \end{aligned} \right]$  Where$\left\{ f\left( x \right)=\left\{ \begin{aligned} 1, &x\geq\varepsilon\\ 0, &\mathrm{otherwise} \end{aligned} \right. \right\}$ |
| Auto Regression coefficient (***AR***) | $\boldsymbol{AR}=\sum_{i=1}^{p} a\left( i \right)x\left( n-1 \right)+e\left( n \right)$  a(i) – autoregressive coefficient  p- autoregressive model order  e(n) - residual white noise |
| Cepstrum Coefficients (***CC***) | -α(i) – $\sum_{i=1}^{i-1} \left( 1- \frac{l}{i} \right)a\left( n \right)c\left( i-1 \right)$  a(i) – autoregressive coefficient  c(i) – Cepstrum coefficient  i – dimensionality of the model |
| Mean Spike Amplitude (*MSA*) [mV] | *SA_i_* = $\frac{\left( Bỿ-Aỿ \right)+(Bỿ-Cỿ)}{2}$  *MSA* = $\sum_{i=1}^{NS} \frac{\mathrm{SA}i}{NS}$  Where Ax, Bx, Cx, Ay, By, and Cy  are the x and y coordinates of the points on the spike |
| Mean Spike Frequency (*MSF*) [Hz] | *MSF* = $\frac{NS}{TD}$ |
| Mean Spike Slope (*MSS*) | *SS_i_* = $\frac{Bỿ-Aỿ}{Bx-Ax}$  *MSS* = $\sum_{i=1}^{NS} \frac{\mathrm{SS}i}{NS}$ |
| Mean Number of Peaks Per Spike (*MNPPS*) | *MNPPS* = $\frac{NP}{NS}$ |
| Mean Spike Duration (*MSD*) [ms] | *MSD* = $\sum_{i=1}^{NS} \frac{Cx-Ax}{NS}$ |
| Modified Mean Absolute Value 1 (*MMAV1*) [mV] | *MMAV1* = $\frac{1}{N}\sum_{i=1}^{N} \left\vert x_{i} \right\vert$  W_i_ = $\left\{ \begin{aligned} 1, if 0.25N\leq i \leq0.75N \\ 0.5, otherwise \end{aligned} \right\}$ |
| Modified Mean Absolute Value 2 (*MMAV2*) [mV] | *MMAV2* = $\frac{1}{N}\sum_{i=1}^{N} \left\vert x_{i} \right\vert$  W_i_ = $\left\{ \begin{aligned} 1, if 0.25N \leq i \leq0.75N \\ \frac{4i}{N}, elseif i<0.25N \\ \frac{4\left( i-N \right)}{N}, otherwise \end{aligned} \right\}$ |
| Simple Square Integral (*SSI*) [mV^2^] | *SSI* = $\sum_{i=1}^{N-1} x_{i}^{2}$ |
| Total Power (*TTP*) [W] | *TTP* = $\sum_{j=1}^{M} P_{j}=SM0$ |
| Power Spectrum Ratio (*PSR*) | *PSR* = $\frac{P_{0}}{P}$ = $\frac{\sum_{j=f_{0}-n}^{f_{0}+n} P_{j}}{\sum_{j=-\infty}^{\infty} P_{j}}$  Where f_0_ is the frequency at which  the maximum Power occurs |
| Higuchi’s Fractal Dimension (*HFD*) | $X_{m}^{k}=\left[ x_{m},x_{m+k},x_{m+2k},{,\ldots,x}_{m+\left\lfloor\frac{N-m}{k} \right\rfloor_{k}} \right]$  Where k is where the integer k is the time  interval between points  and $\left\lfloor\frac{N-m}{k} \right\rfloor_{k}$is the integer part of $\frac{N-m}{k}$  $L_{m}= \frac{1}{k}\left[ \frac{N-1}{\left\lfloor\frac{N-m}{k} \right\rfloor k} X\sum_{i=1}^{\left\lfloor\frac{N-m}{k} \right\rfloor} \left\vert x_{m+ik}- x_{m+\left( i-1 \right)k} \right\vert\right]$  L(k)=$\frac{\sum_{m=1}^{k} L_{m}(k)}{k}$  L(k) the length of the curve for time interval k |
| Maximum fractal length (*MFL*) | *MFL*=$\log_{10} \left( \sqrt{\sum_{i=1}^{N-1} \left( x_{i+1}-x_{i} \right)^{2}} \right)$ |
| Skewness (*SKEW*) | *SKEW* = ($\frac{1}{N}\sum_{i=1}^{N} (\frac{x_{i}- x͞}{\sigma}$)₃) |
| Kurtosis (*KURT*) | *KURT* =($\frac{1}{N} \sum_{i=1}^{N} (\frac{x_{i}-x͞}{\sigma})₄)-3$ |
| Average Amplitude Change (*AAC*) [mV] | $ACC={\frac{1}{N}}\sum_{i = 1}^{N -1} \vert x_{i+1}x_{i} \vert$ |
| Difference Absolute Standard Deviation Value (*DASDV*) [mV] | $DASD={\sqrt{\frac{1}{N-1} \sum_{i=1}^{N-1} (x_{i+1 -} {x_{i})}^{2}}}$ |
| Myopulse Percentage Rate (*MYOP*) | $MYOP={\frac{1}{N}}\sum_{i = 1}^{N} [ f\left( xi \right) ]$  Where $\left\{ f\left( x \right)=\left\{ \begin{aligned} 1, &x\geq\varepsilon\\ 0, &\mathrm{otherwise} \end{aligned} \right. \right\}$ |
| Mean Absolute Value Slope  (*MAVS*) [mV] | *MAVS*= ${MAV}_{k+1}$ - ${MAV}_{k}$ |
| Multiple Window (***MW***) [mV] | $\boldsymbol{MW}_{\mathbf{k}}$=$\sum_{i=0}^{N-1} \left( x_{i}W_{i-i_{k}} \right)^{2}$ , k=1,…,K  Where W_i_ is the i^th^ value of the windowing function |
| Shannon entropy (*SHANNON*) | *SHANNON*=-$\sum_{j=1}^{M} p_{j}\ln(p_{j})$  Where p_j_ is the probability of the j^th^ outcome |
