## Appendix 2 for "Modeling the assessment of the upper limb motor function impairment in children with cerebral palsy using sEMG and IMU sensors"

Individual performance for each feature with majority vote among all tasks (ranked in order of RF classification accuracy for sEMG and LDA for IMU-ACC)

| **Feature** | **LDA** | **SVM** | **RF** |
| --- | --- | --- | --- |
| **sEMG features** | | | |
| mss | 71.43 | 53.57 | 85.71 |
| mw | 50.00 | 53.57 | 82.14 |
| aac | 67.86 | 53.57 | 82.14 |
| iemg | 64.29 | 42.86 | 82.14 |
| sm2 | 64.29 | 67.86 | 82.14 |
| mtw | 57.14 | 53.57 | 78.57 |
| msa | 71.43 | 53.57 | 78.57 |
| dasdv | 75.00 | 53.57 | 78.57 |
| log | 67.86 | 53.57 | 78.57 |
| mav | 60.71 | 53.57 | 78.57 |
| mdf | 71.43 | 42.86 | 78.57 |
| mfl | 60.71 | 67.86 | 78.57 |
| mmav2 | 64.29 | 53.57 | 78.57 |
| pkf | 78.57 | 46.43 | 78.57 |
| sm1 | 57.14 | 53.57 | 78.57 |
| sm3 | 71.43 | 46.43 | 78.57 |
| mmav1 | 60.71 | 53.57 | 75.00 |
| mnp | 71.43 | 53.57 | 75.00 |
| ttp | 60.71 | 53.57 | 75.00 |
| wl | 71.43 | 64.29 | 75.00 |
| hfd | 78.57 | 75.00 | 71.43 |
| mavs | 60.71 | 53.57 | 71.43 |
| MNF | 67.86 | 46.43 | 71.43 |
| rms | 75.00 | 53.57 | 71.43 |
| ssi | 71.43 | 53.57 | 71.43 |
| var | 71.43 | 53.57 | 71.43 |
| mnpps | 71.43 | 71.43 | 67.86 |
| cc4 | 60.71 | 39.29 | 67.86 |
| fr | 71.43 | 67.86 | 67.86 |
| psr | 71.43 | 67.86 | 67.86 |
| zc | 64.29 | 46.43 | 67.86 |
| msf | 67.86 | 46.43 | 60.71 |
| msd | 75.00 | 53.57 | 60.71 |
| shanNon | 42.86 | 57.14 | 60.71 |
| skew | 64.29 | 42.86 | 60.71 |
| ssc | 50.00 | 46.43 | 60.71 |
| kurt | 50.00 | 46.43 | 57.14 |
| myop | 57.14 | 53.57 | 53.57 |
| vcf | 57.14 | 46.43 | 53.57 |
| AR4 | 39.29 | 42.86 | 50.00 |
| wamp | 53.57 | 46.43 | 46.43 |
| **Accelerometric features** | | | |
| Mean | 89.26 | 85.71 | 85.71 |
| Maximum | 78.57 | 89.29 | 78.57 |
| Minimum | 75.00 | 89.26 | 82.14 |
| Minima | 75.00 | 46.43 | 67.86 |
| Maxima | 75.00 | 46.43 | 67.86 |
| Standard deviation | 46.43 | 57.14 | 57.14 |
